# Male behavior in a swallowtail butterfly (*Battus philenor*) ensures directional iridescent sexual signal is visible to females during courtship

**DOI:** 10.1101/2023.04.03.535474

**Authors:** Ronald L. Rutowski, Nicolas Lessios, Brett Seymoure, Kim Pegram, Andrew Raymundo

**Affiliations:** School of Life Sciences, Arizona State University, Tempe, AZ, USA 85287-4501; Department of Biological and Physical Sciences, Assumption University Worcester, MA, USA 01609; Department of Biological Sciences, University of Texas at El Paso, El Paso, TX, USA 79968; Desert Botanical Garden, 1201 N. Galvin Parkway, Phoenix, AZ USA 85008

## Abstract

Iridescent colors in animals may vary with the arrangement of the observer and the light source relative to the animal. When these highly directional colors are used as signals, there may be transmission problems for senders in that the perception of the signal by intended receivers can be greatly affected by the spatial arrangement of the sender, receiver, and sources of contributing light. One potential solution to this problem is for senders to behave in a manner that arranges the positions of sender, light source, and receiver to enhance signal perception by the receiver. We tested this idea by describing the structure of courtship behavior of males of the Pipevine Swallowtail (*Battus philenor*) butterfly and analyzing its consequences for the female detection of the male’s iridescent sexual signal which is used in mate choice by females. During courtship, males perform a swoop maneuver that brings their dorsal hindwing from below to directly in front of the female. Several swoops in rapid succession occur in aerial courtships that lead to copulation. We measured under solar radiation the radiance of the male’s dorsal hindwing as seen by the female during a swoop. Regardless of how the male’s swoop path is oriented relative to the sun, there will be a brief, bright, and saturated flash of blue visible to the female. Our results support the conclusion that male swoops enhance the visibility of the male’s iridescent color signal to the female during courtship.

## INTRODUCTION

As predicted by the sensory drive hypothesis (Endler 1992, Endler and Basolo 1998), male animals often behave during courtship in ways that enhance the transmission of signals used in mate selection. For example, in the Queen butterfly (*Danaus gilippus*), male behavior during courtship, called hairpenciling, brings their scent-producing structures into close proximity with the female’s antennal chemosensors (Brower et al. 1965). For directional color signals, the courtship behavior of males is expected to include features that adaptively enhance the visibility of their iridescent signals to females, as was first suggested by Poulton (1890).

Many iridescent colorations are mirror-like or specular which mean they are only visible in specific spatial arrangements of the light source (typically the sun), the reflecting surface, and the viewer. Moreover, it is not just the brightness that will vary with this spatial arrangement. As brightness changes so will the saturation of the visible color change. Iridescent colors by definition vary in hue with view angle (Simon 1971), which might change during courtship. This all means that in species in which females make mate choice decisions based on male iridescent color signals, the relative positions of the male, female, and sun during courtship will importantly determine signal visibility and reliability (Stuart-Fox et al. 2021, Doucet and Meadows 2009).

In a few butterfly species, males bear iridescent patches on their wings that have been inferred or demonstrated to be sexual signals used by females in their selection of mates (Kemp and Rutowski 2011; Rutowski and Rajyaguru 2013). In our studies of the iridescent sexual signal of the male of the Pipevine Swallowtail (*Battus philenor*; Fig. 1), we (Rutowski et al. 1989, Rutowski et al. 2010) and others (Mitra et al. 2016) have observed behavior in males that appears to fit the expectation that males are enhancing the visibility of their directional color signal to females. Specifically, during courtship, a male flies from below a flying female up in front of her, then moves back and over her before he drops below and behind the female before repeating the maneuver. Mitra et al. (2016) called this circular maneuver a swoop and we will follow and formalize that terminology here.

**Fig. 1.**
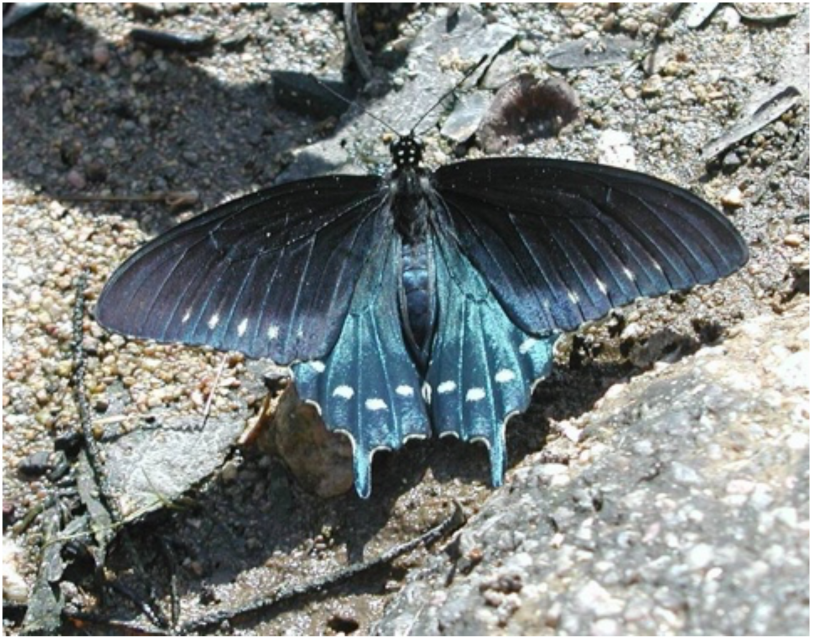
Dorsal wing surfaces of male Pipevine Swallowtail, *Battus philenor*. The blue reflections are used by females in mate choice and are iridescent.

Here we report detailed descriptions and analyses of males’ swoops during courtship using high frame rate videography. We begin with a general description of courtship leading to mating in the Pipevine Swallowtail and then describe quantitatively from the video recordings how the male positions himself relative to the female during swoops. Finally, we present analyses of the saturation, brightness, and hue of the radiance from the male’s wings that would appear to a female under natural lighting during his upward movement during a swoop. Our data are intended to test the prediction that the swoops are structured in a way that makes his directional blue signal maximally visible to the female.

Our study parallels both in purpose and overall design a number of recent studies evaluating the appearance to receivers of iridescent colorations during behavioral interactions in nature (e.g. Sicsú et al. 2013, Schultz and Fincke 2009, Simpson and McGraw 2018a, 2018b, 2019a), but especially White et al.’s (2015) innovative study of the courtship behavior of males of the Common Eggfly, *Hypolimnas bolina*. That study evaluated how the male’s distinctive courtship flight pattern affects the visibility of his iridescent signal to the target female. Our study complements and extends these studies in two ways. First, we describe in previously unreported detail a courtship display that is very different from that of *H. bolina* and so represents a potential novel adaptive solution to the problem of making an iridescent signal visible to a potential mate. In the discussion, we compare our observations of courtship in *B. philenor* to what is known more generally about interspecific variation in the behavior of male butterflies leading to copulation to explore the evolutionary novelty of the swoop. We also evaluate the generality of statements about butterfly courtship behavior made by Darwin in 1871.

Second, we attempt to characterize the visibility of the male’s iridescent signal to females during courtship under conditions that closely match those in nature, in much the same way White et al. (2015) did but using a very different protocol. Instead of inferring male appearance to females from measures of male wing surface reflectance and of ambient irradiance, we measured the radiance directly from the male’s wings in the field under specific conditions chosen from our understanding of the natural history of *B. philenor*. First, we focused our measurement on morning hours. Females eclose around sunrise and take their first flight and are most likely to be detected and courted by males in the morning (Hannam et al., 2018). Females are receptive at this time (Rutowski et al. 1989; this study). Second, we made our measurements when the sun was at solar elevations typical of spring and fall days (around 50°; Rutowski et al. 1989) when these butterflies are seasonally most active, at least in the Sonoran Desert. Finally, we selected other conditions for the measurement of radiance described in the Methods that were based on the details of the males’ behavior during courtship that we document in our study. Our strategy for evaluating the appearance of the male’s radiance during swoops under natural light, follows the recommendation of Fleishman et al. (2006) for how best to assess the appearance of specular colorations in nature.

Again, our overall prediction was that the courtship display of the male will enhance the visibility of his blue iridescent signal to the female in nature, at least at some point or points during the display.

## METHODS

### Origins of animals studied

All observations were made on adult butterflies that were reared from eggs and larvae collected in the vicinity of Mesquite Wash (33.7293°N, 111.5117°W) in the Mazatzal Mountains in central Arizona. At this location, *Aristolochia watsonii*, the larval foodplant of the Pipevine Swallowtail is common. We collected tendrils of this plant and fed them to larvae housed in the lab where they were kept at 16hrs of light at 30°C, 8hrs of dark at 24°C, and a constant 55% relative humidity.

### Courtship observations, recordings, and analysis

Observations of courtships leading to mating were made on animals placed within the Maxine and Jonathan Marshall Butterfly Pavilion at the Desert Botanical Garden in Phoenix Arizona described previously (e.g., Rutowski and Rajyaguru 2013). We populated the pavilion with up to 10 lab-reared males and then presented them with lab-reared virgin females either by placing a female on a conspicuous perch and coaxing her into flight as a male flew by or by releasing a female into the flight path of a male. Ensuing courtships were observed and each described as to whether the female was initially perched or flying when the male arrived within 10 cm of her, whether the pair mated, the number of swoops performed by the male, and the total duration of the interaction. Durations were timed to the nearest 0.1 sec with a handheld stopwatch from when the male first arrived within 10 cm of the female until he either succeeded in coupling with the female or departed and did not return within a few seconds. In some cases, pairs were quickly and gently separated and the female observed in up to as many as three subsequent courtships leading to copulation. This was done with only 11 of a total of 22 females. In three cases, the same male was observed in 2-3 courtships.

The speed, three-dimensional structure, and brevity of successful courtships precluded success in capturing them in video recordings. However, some unsuccessful courtships were lengthy and allowed us to capture them digitally. Video recordings were made using a handheld Casio Exlim ZR100 camera at 240 frames/sec. For analysis of the position of the male relative to the female during courtship, we selected 17 clips in which both the female and courting male were shown from the lateral perspective (see example in Supplemental Materials). This provided a good view of the swoop path which tended to be in the same plane as that defined by the female’s dorso-ventral and anterio-posterior axes. Starting with the male below and behind the female, at intervals of 10 frames, we laid a transparent X-Y grid over the screen with the origin on the female’s head. We then recorded the X-Y coordinates of location of the male’s thorax in the frame. The coordinates at each time interval were converted to units of female forewing length using the length of the female’s forewing in grid units. The sample size at each time interval shown in Fig. 3 varies due to variation in clip lengths and because position of the male’s thorax and female’s head were not clear in every frame due to motion blur, shadows, and positioning. These data allowed a detailed description of the time course of the changes in the male’s position relative to the female during a swoop.

From these video clips we were also able to describe the orientation of the male swoop path and the female flight path relative to each other and the sun. Path bearings were determined to the nearest 45° using the known position and compass alignment of landmarks in the pavilion that were visible in the clips, including the walls of the enclosure and plants and objects visible inside and outside the enclosure. Solar positions for this analysis were obtained from the NOAA Solar Position calculator (https://gml.noaa.gov/grad/solcalc/azel.html).

### Radiance measurements

To evaluate the appearance of a male to a female during a swoop we measured the dorsal hindwing radiance under natural illumination. To measure radiance, we used a collimating lens affixed to an optical fiber that was attached to an Ocean Optics USB 2000+ spectroradiometer. The spectrometer was connected to a laptop running SpectraSuite software (Ocean Optics, Dunedin, FL). The fiber without the lens was calibrated for radiance measurements using an Ocean Optics LS CAL calibrated light source. With the lens added, the system reported light input in microwatts/cm^2^/sec. For color analysis, we converted this to radiance units of photons/cm^2^/sec/sr, taking into consideration the collimating lens was collecting light from a solid angle of 0.00165 sr.

The measurements were made between 9:00 and 11:00 on cloudless days (15 May, 16 May, and 6 July 2014) in the middle of a large plaza on the ASU campus (33.4205°N, 111.9340°W). At this time of day on these dates the solar elevation varied from about 45 to 65° above the horizon. Changes in solar azimuth were tracked as described below. This time of day was chosen because 1) females eclose from the pupa around sunrise and spend an hour or more expanding and hardening their wings (Hannam et al. 2018), and 2) males begin flying around 7:00 (Pegram et al. 2012). From this, we infer that most courtships begin around midmorning with males approaching virgin females that have begun to fly.

A more detailed analysis is presented in the results, but we determined from the above analysis that during a swoop, the male flies from directly below the female in an upward arc that brings him in front of the female with his body axis vertical as he passes in front of the female. This arc is in the same plane as the flight path of the female. Hence during a swoop, the pitch of a male’s wing plane relative to the horizontal plane goes from about 0° when flying level below the female to 90° as he passes in front of the female (Fig. 2). Our aim was to determine at what wing pitches during a swoop the peak of the spectrum as measured from the position of the female would be at a maximum as seen on the laptop. We collected the spectrum of the radiance at this maximum to determine how the wing pitch at which this peak occurs varied with the orientation of the male’s wing relative to the solar azimuth.

**Fig. 2.**
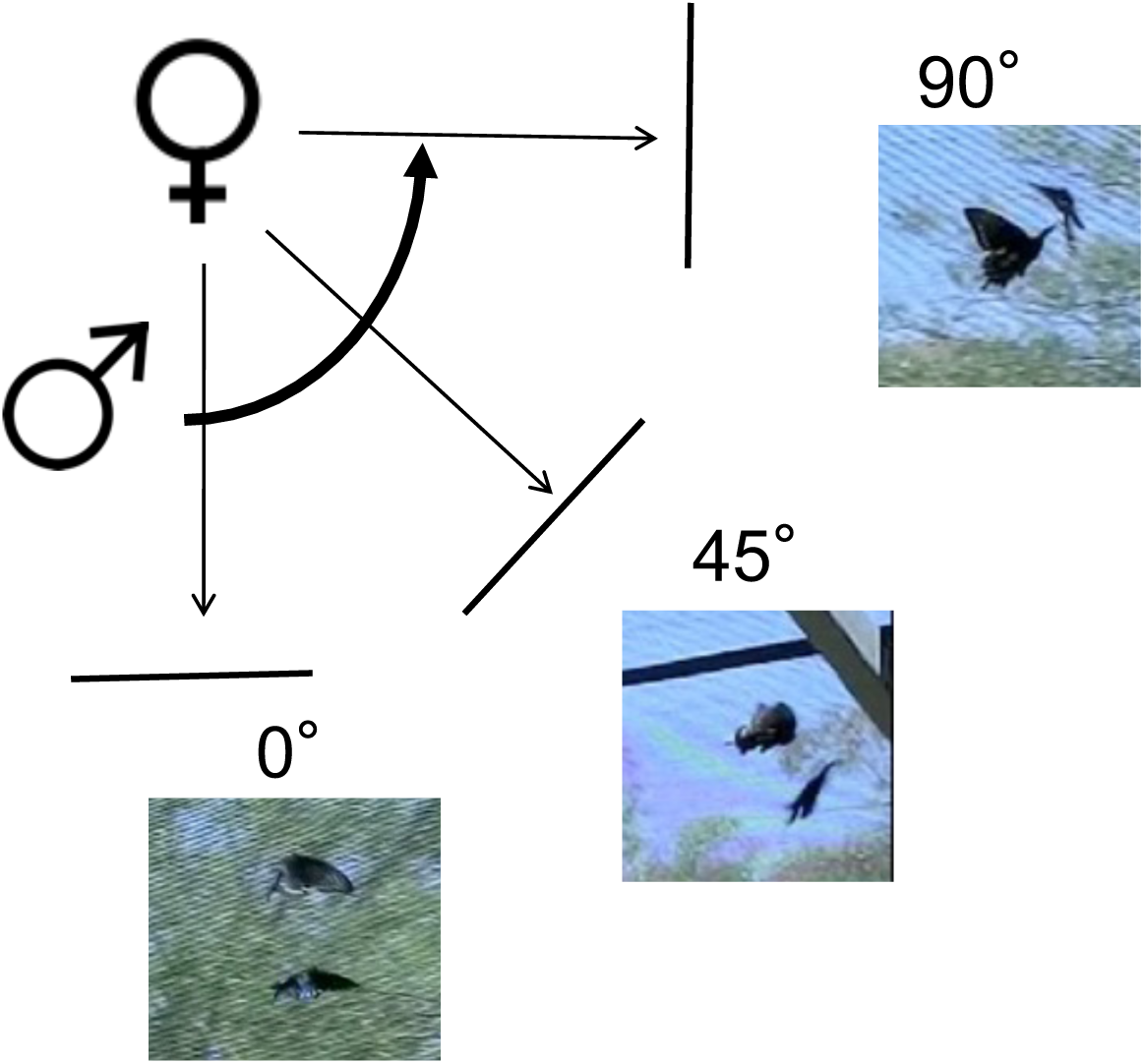
A diagram showing the path of a male relative to a female during a swoop. The photographs show how the wing pitch of the male increases from 0° relative to horizontal at the beginning of swoop to 90° when the male come up directly in front of the female. In each image, the female is the individual above or to the left of the other individual. For radiance measurements, the collector was placed in the position of the female.

We placed a male hindwing on the stage of a custom-made light table that allowed us to manipulate the pitch and the orientation of the wing relative to the solar azimuth (Meadows et al. 2011). We pointed the long axis of the wing (wing base to tail) so that the line from the tail to the wing base pointed at each of eight angles relative to the solar azimuth (0°, 45°, 90°, 135°, 180°, 225°, 270°, and 315°). At each of these orientations we changed the wing pitch through 10° increments keeping the axis of the collector light path normal (perpendicular) to the wing surface (Fig. 2) until again, the peak of the radiance spectrum monitored on the laptop was at maximum. We collected the spectrum at this maximum for color analysis and recorded the wing pitch.

For color analysis, we used the software CLR (Montgomerie 2008). With this software we truncated collected radiance spectra to 300-700 nm, binned the data into 10 nm intervals, and calculated three color parameters. Brightness was the calculated mean of the radiance in the measured spectrum (B2 in CLR). In previous studies of reflectance of the iridescent blue in *B. philenor* relative to a white standard (e.g., Rutowski et al. 2010), we used chroma S3 (a measure of spectral purity) and hue H1 (a measure of spectral location) calculated by CLR. Both of these measures rely on determining the wavelength of peak reflectance, λ_Rmax_, However, in this study of radiances, these were inappropriate to use. Because the spectrum of sunlight is not a flat smooth curve - it has many small peaks and valleys (Endler 1993) −, the radiances we measured were also not smooth curves. Under these conditions, the wavelength of peak reflectance, λ_Rmax_, in the radiance spectra can be difficult to pinpoint.

Hence, for this radiance study, we used this measure, which we have here for clarity called saturation (S5c in CLR, normalized to the total brightness).

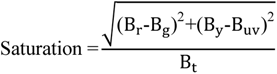

For hue, we used this hue angle measure which, again, does not depend on λ_Rmax_.

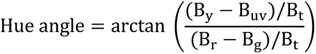

In both equations, the subscripts are as follows: r = 600-700 nm, y = 500-600 nm, g = 400-500 nm, uv = 300-400 nm, t = 300-700nm. B_x_ is the summed brightness in each of these spectrum segments, respectively.

Hindwings from six males were used for measurements of radiance brightness, saturation, and hue angle that were analyzed in Tests A1-A4 (Table 1). All males were from the control group (dorsal hindwing coloration not altered) in the mate choice experiment described in Rutowski and Rajyaguru (2013). This sample of six males was chosen to evaluate the appearance during a swoop of males across a range of chroma values which were significantly higher in unaltered mated males in Rutowski and Rajaguru (2013). Three were males from among those that had mated and from reflectance measurements in the lab had the highest chroma (S3; mean ± SD: 0.469 ± 0.014; range: 0.461-0.4485), and three were from among those that had not mated with the lowest chroma of their reflectance (S3; mean ± SD: 0.40 ± 0.015, range: 0.38-0.409; t-test, p<0.05). In this sample of males, the mean brightness and hue of their reflectance did not differ (t-test, p>0.05) between the two groups (brightness (mean ± SD): mated = 12.1% ± 1.74%; range: 11.5-14%, unmated = 11.1% ± 3.93%; range: 11.5-14%; λ_Rmax:_ (mean ± SD): mated = 489.2 ± 5.48 nm, unmated = 463.5 ± 20.1 nm).

**TABLE 1:** Results of ANOVAs analyzing how radiance properties are affected by swoop direction relative to the solar azimuth (Tests A1-A4; Two-Way Repeated Measures ANOVA) and wing pitch relative to the pitch at maximum brightness (Tests B1-B3; One-Way Repeated Measures ANOVA). Group refers to whether males were mated or not in a prior experiment. Significant effects (p<0.05) are shown in **bold**. See text for details.

| Test | Dependent Variable | Repeated Measure | Within Subjects Effects | Between Subjects Effects |
| --- | --- | --- | --- | --- |
| A1 | Swoop direction | Wing pitch at maximum radiance | **Swoop dir.: $F_{1,7}=28.7, p<0.001$** | Group: $F=0.4.95, p=0.521$ |
| | | | Swoop dir.*Group: $F_{1,7}=1.25, p=0.311$ | |
| A2 | | Brightness | Swoop dir.: $F_{1,7}=0.672, p=0.458$ | Group: $F=0.112, p=0.755$ |
| | | | Swoop dir.*Group: $F_{1,7}=0.424, p=0.55$ | |
| A3 | | Hue angle | Swoop dir.: $F_{1,7}=3.52, p=0.134$ | Group: $F=21.2, p=0.01$ |
| | | | Swoop dir.*Group: $F_{1,7}=3.64, p=0.129$ | |
| A4 | | Saturation | Swoop dir.: $F_{1,7}=7.06, p<0.001$ | Group: $F=0.347, p=0.587$ |
| | | | Swoop dir.*Group: $F_{1,7}=3.06, p=0.016$ | |
| B1 | Wing pitch relative to pitch at maximum radiance | Brightness | **Rel. Pitch: $F_{1,6}=33.1, p<0.001$** |  |
| B2 |  | Hue angle | **Rel. Pitch: $F_{1,6}=25.7, p<0.001$** |  |
| B3 |  | Saturation | **Rel. Pitch: $F_{1,6}=17.8, p<0.001$** |  |

However, when the reflectance spectra of these males were analyzed using the saturation and hue angle measures used in this study, mated males had a significantly higher (bluer) hue angle (mean ± SD: mated = 2.72 ± 0.24 radians, unmated = 2.04 ± 0.08 radians, t-test, p < 0.05) but did not differ in saturation (mean ± SD: mated = 2016 ± 382, unmated = 1505 ± 671, t-test, p > 0.05). We therefore entered the male’s group (mated vs. unmated) in our analyses to see how the radiance color parameters at maximum radiance changed with orientation of the swoop relative to the solar azimuth in the field and how these differed among male groups.

A second set of measurements were obtained from a set of four male hindwings selected haphazardly from lab-reared specimens used in Rutowski et al. (2010). Tests B1-B3 in Table 1 present the analysis of these measurements which were intended to reveal how the color of the male changed during the performance of a single swoop. In this case we examined how the color parameters changed as the wing pitch was set at −30°, −20°, −10°, 0°, 10°, 20°, and 30° relative to the wing pitch that produced the highest brightness. As the hindwings were selected haphazardly, whether the male was mated or not was not relevant to our analysis. All other aspects of the protocol were identical to those already described.

### Statistics

Descriptive and inferential statistics described in the Results for continuous variables were calculated for t-tests with Microsoft Excel, and for ANOVAs, IBM SPSS 28. All color parameter data sets that were analyzed with a repeated-measures ANOVA were tested and satisfied the assumption of normality. Flight path bearings for males and females during swoop were measured in degrees as described and summarized using descriptive circular statistics. Bearing distributions were compared among data sets evaluated with Rayleigh tests of uniformity. For this, we used the software package Oriana (Kovach, 2011). All inferential test statistics were evaluated at the 0.05 level of significance.

## RESULTS

### General description of successful courtship

Courtships leading to copulation are so brief (see below) and occur so unpredictably in space and time, that filming those interactions at adequate spatial and temporal resolution for detailed analysis was not feasible. The qualitative description that follows is from direct observations with the naked eye.

In courtships leading to copulation that begin with the female flying, the male typically approaches the female and takes up a position below and behind her. He often matches the female’s forward velocity briefly which causes him to hold this position below the female with the long axis of his body roughly parallel to the ground and with the same bearing as that of the female. Next, he accelerates upward and forward relative to the female which brings him up in front of the female. During this acceleration, his body axis becomes increasingly vertical so that as he passes upward in front of the female his body axis is vertical and his wings are held open in front of the female. Following Mitra et al. 2015, we call this action a swoop (see video in Supplementary Material). During this male maneuver the female continues to fly steadily forward.

Once the male has started to move upward from a position directly in front of the female to a position above her, his forward velocity drops so that the female moves ahead of and underneath him. His body axis returns to horizontal as he drops back to a position again below and a bit behind the female. He then accelerates and performs another swoop, often performing several in rapid succession. After a series of swoops the female alights on vegetation with the wings closed at which point the male alights alongside her facing the same direction and brings the tip of his abdomen into contact with hers.

We timed 24 courtships leading to mating with this general form and their durations averaged 17.2 ± 1.9 sec (range = 6.1-45 sec). Our counts of the number of swoops in 20 successful courtships (female initially flying) yielded an average of 9.7 ± 1.7 swoops/courtship (range = 1 - 38 swoops/courtship). This suggests a swoop rate of about two swoops per second. All successful courtships with females that were initially flying included swoops by the male.

We observed fewer successful courtships in which the female was perched when first approached and remained so until mating occurred (n = 9). The duration of these courtships averaged 7.97 ± 1.81 sec (range = 4.6 – 22 sec). In these courtships leading to copulation, males did not perform swoops which indicates that, in some circumstances, copulation can occur without them.

### Swoop structure

In unsuccessful courtships, flying females approached by males often do not alight but keep flying as the male performs repeated swoops. The longer duration of these courtship allowed us more readily and quickly to position the camera for high frame rate videography. The video recordings obtained permitted a detailed description of the spatial and temporal structure of the male’s behavior during a swoop, albeit during unsuccessful courtships. From 29 clips that showed consecutive swoops, we calculated the time from when the male was directly in front of the female in the first swoop until he was directly in front of her in the second swoop. The inverse of this time was the swoop rate which averaged 1.61 ± 0.083 swoops/sec (range = 0.7 – 3.07 swoops/sec). This agrees generally with the swoop rate estimated from the counts of the number of swoops in successful courtships.

We obtained high-frame-rate clips showing 17 swoops viewed laterally, that is, taken from the same height as the female above the ground and perpendicular to her flight path. Because no size reference could be placed in the frame as these clips were made, we made all linear measurements in units of female forewing length. For a given clip this was measured when the female’s wings were closed over her back during a wingbeat cycle. In a previous study the average forewing length of females reared in our lab like those in this study was 5.18 cm (Rutowski et al. 2010).

From these 17 clips, we analyzed the movements of the male relative to the female during a swoop in detail. The results shown in Fig. 3 and made during the course of these measurements allow several statements about how the males are behaving. First, during a swoop the male’s point of closest approach to the female is when he is directly in front of her. At this point he is within one forewing length (about 5 cm) of the female. Second, the male initiates his flight up and in front of the female about 175 msec before he is directly in front of the female. At the beginning, he is about 2 forewing lengths (about 10 cm) below the female. Third, his path during the upward phase of the swoop is much less variable than at other points in the swoop cycle in both the horizontal and vertical directions. This suggests that the upward part of the swoop is a highly ritualized or stereotyped flight movement. Fourth, during the upward movement of the swoop we observed that the male’s wings are held open and not flapped. Fifth, although not measured directly, during the upward phase of the swoop the male’s body pitch increased from about 0° to about 90° relative to horizontal as he passes directly in front of the female. As a result, there is a corresponding change in the pitch of the wings from about 0° to about 90° relative to horizontal (Fig. 2).

**Fig. 3.**
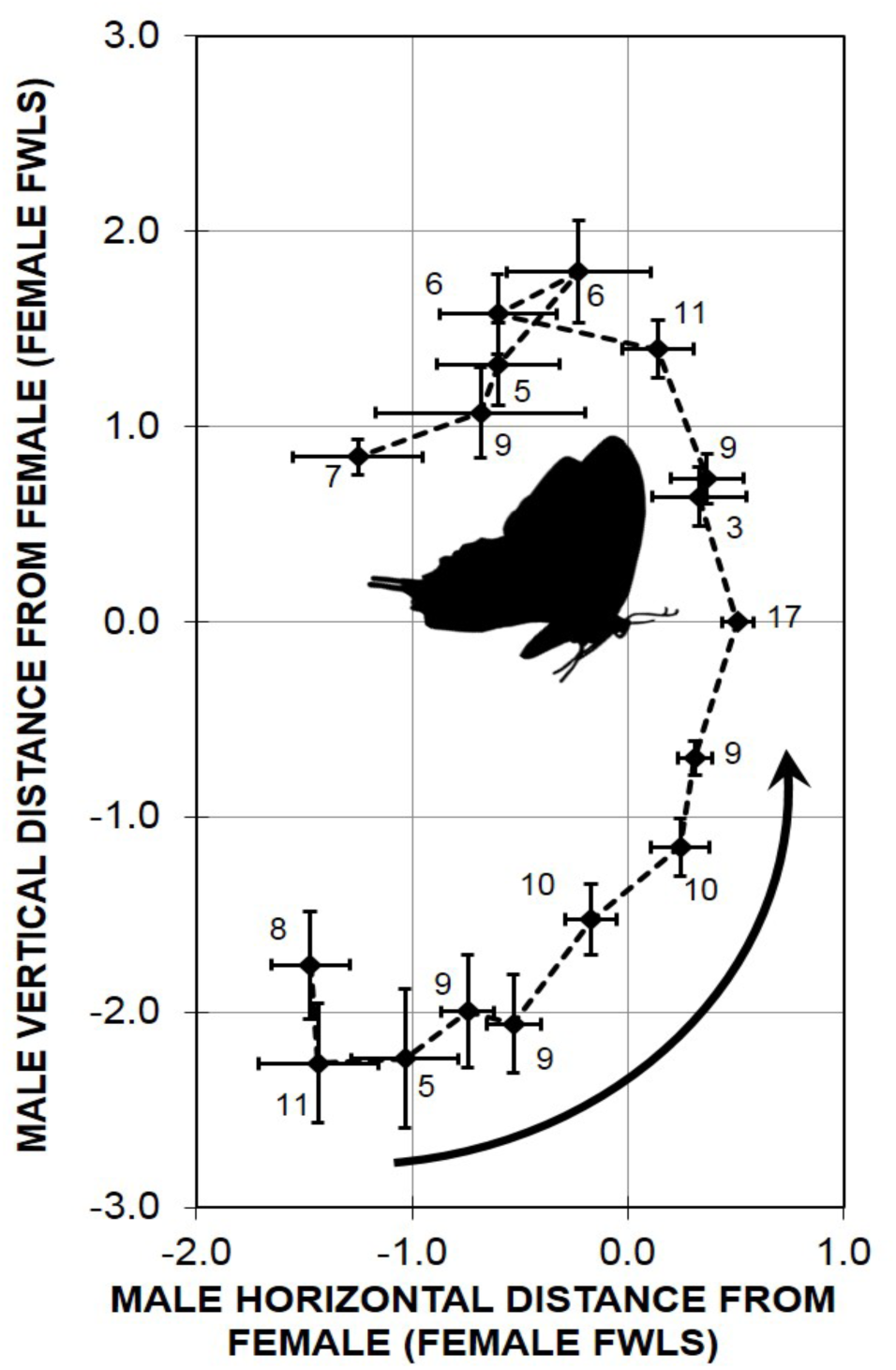
The path taken by a male *B. philenor* relative to the flying female’s head (shown in silhouette) during a single swoop starting from under the female and ending above the female. The plot summarizes data from 17 swoops. The mean position of the male, the standard error of the mean in the vertical and horizontal direction, and the sample size are shown at 25 msec intervals from the time before and after the male is directly in front of the female.

### Swoop path relative to sun

Several results from our video recordings of swoops in unsuccessful courtships indicate that the flight bearing of a female being courted is the main determinant of the bearing of swoops performed by the courting male. The flight paths of females at the time of a swoop were oriented randomly relative to north (Fig. 4). Moreover, the bearings of swoops were oriented randomly relative to the solar azimuth (Fig. 5) but were very much aligned with the flight path of a female being courted (Fig. 6).

**Fig. 4.**
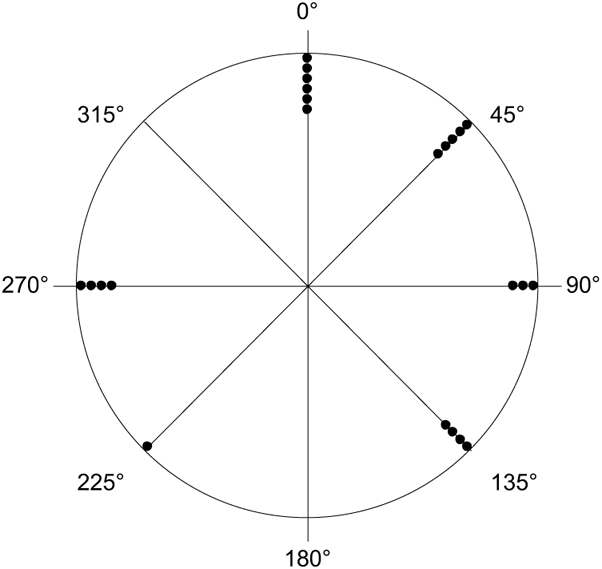
The flight bearing relative to North of females (n=23) during a swoop directed at them. Females being courted are not flying any consistent direction relative to magnetic north. Rayleigh test of uniformity: p = 0.07.

**Fig. 5.**
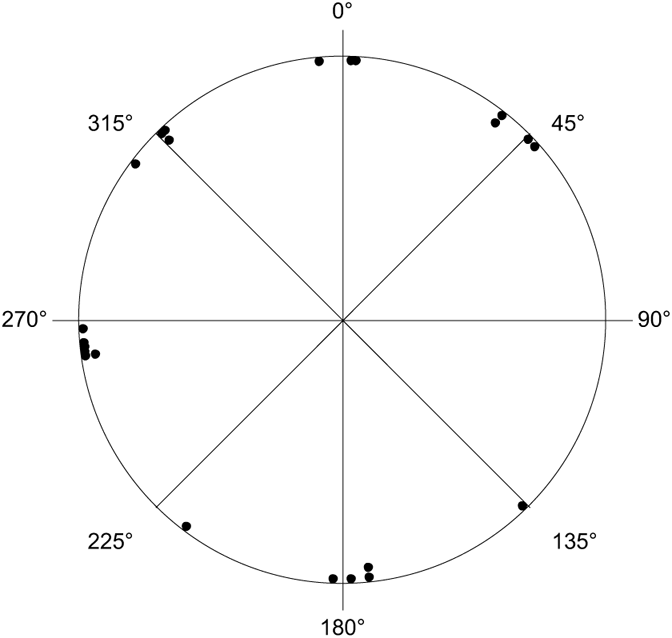
The difference between the bearing of a swoop and the bearing to the solar azimuth (n = 23). There is no consistent relationship between the swoop bearing and the bearing to the solar azimuth. Rayleigh test of uniformity: p = 0.172.

**Fig. 6.**
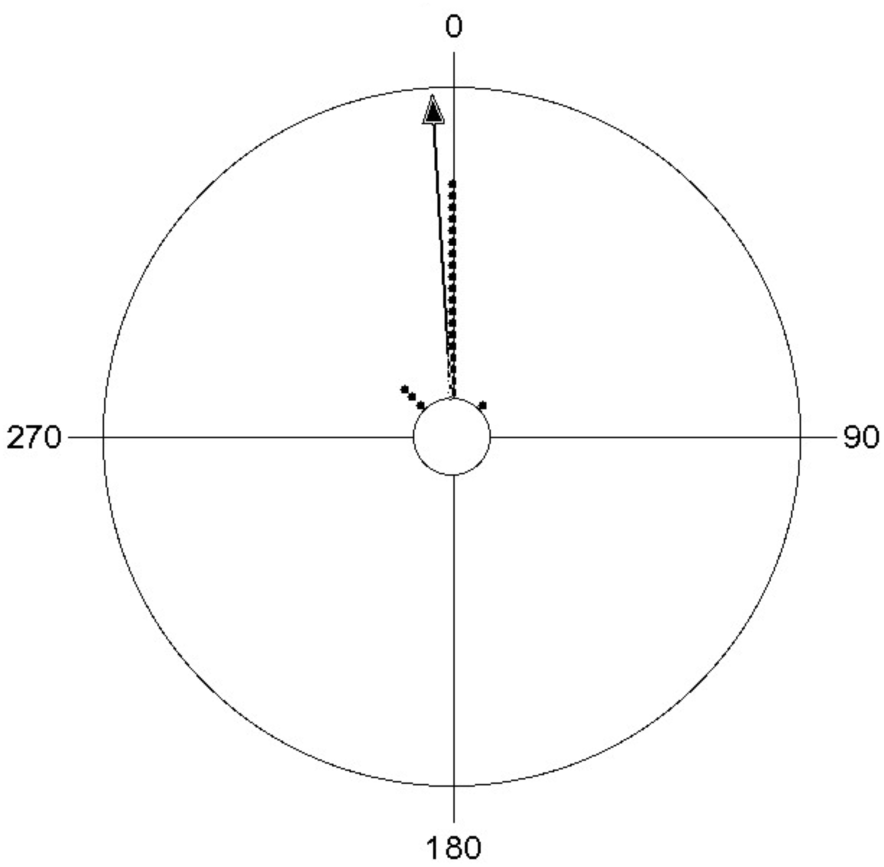
The difference between the bearing of a swoop and the female flight bearing (n = 23). Swoop bearing generally matches that of the target females flight path. Rayleigh test of uniformity: p < 10^-8^, mean vector = 356⁰, vector length = 0.975.

### Dorsal hindwing radiance during a swoop from the female’s perspective

We evaluated how the orientation of a swoop path relative to the sun’s azimuth affected female perception of the iridescent coloration on the male’s dorsal hindwing surface as wing pitch changes during the upward phase of a swoop. To approximate the direction of the swoop path relative to the sun’s azimuth in each measurement, we used the direction the hindwing tail was pointed relative to the solar azimuth. For example, if the tail was pointing toward the azimuth, for that measurement the swoop direction was described as 180 degrees relative the solar azimuth.

The wing pitch at which dorsal hindwing radiance was at a maximum varied with the orientation of the swoop path relative to the solar azimuth (Fig. 7; Table 1, Test A1, Swoop dir.). Specifically, when the swoop path was away from the solar azimuth (bearing 180°; sun behind the male), the wing pitch yielding maximum radiance was at or a little below 0°, basically horizontal as it is at the beginning of a swoop. However, when the swoop path is toward the sun (bearing 0°), the wing pitch of maximum radiance is about 60° relative to horizontal, as it is when the male is approaching being directly in front of the female. These pitch angles are like those that the wings will pass through during a swoop (0 to 90°). Across swoop directions, the pitch angle of maximum radiance for the mated males was not different from that for unmated males (Table 1, Test A1, swoop direction*group).

**Fig. 7.**
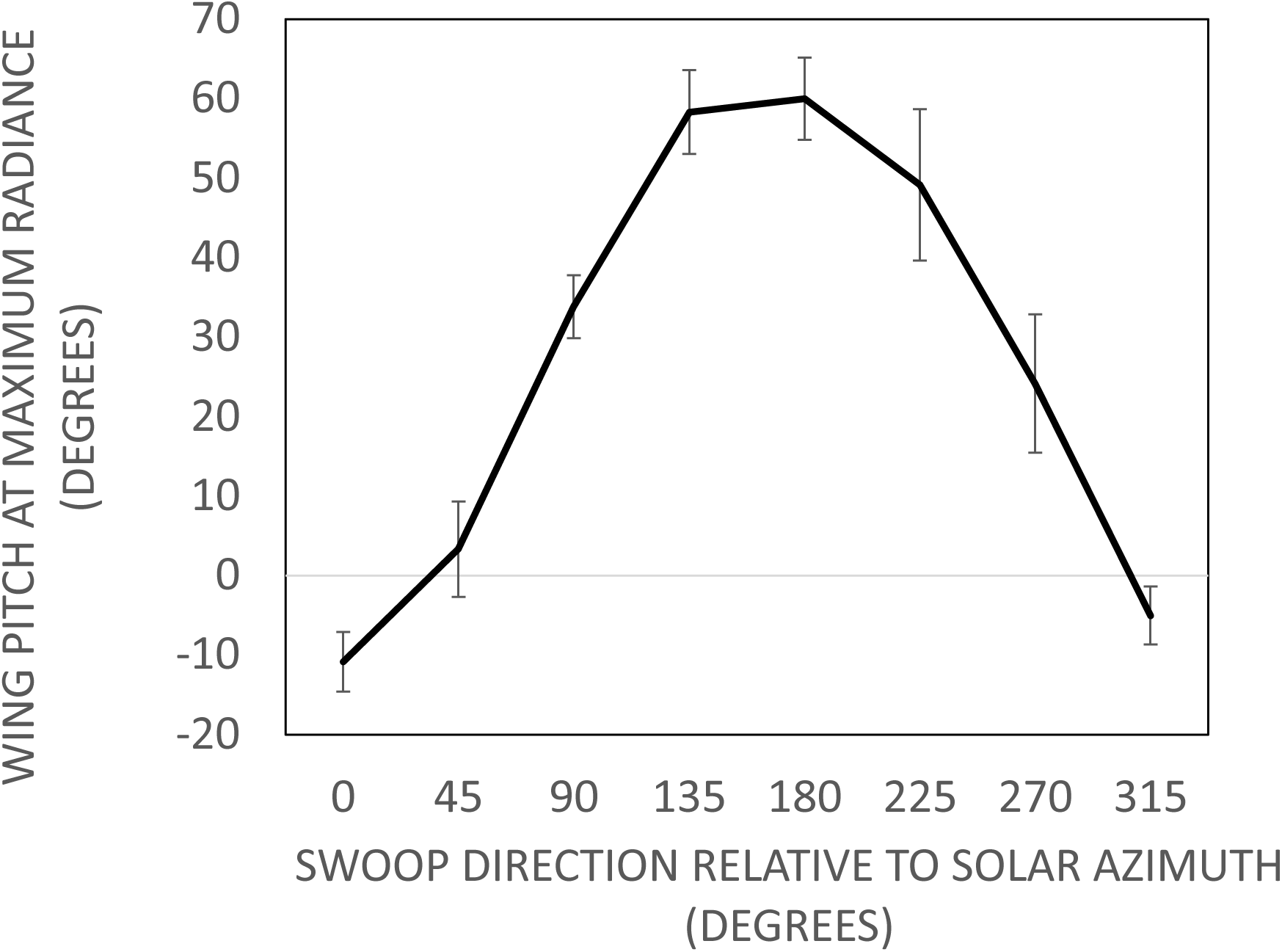
How the orientation of the path of a male during a swoop relative to the sun affects when during a swoop his iridescent blue coloration will display the greatest radiance from the perspective the female being courted. Six male butterflies were sampled. The error bars indicate the mean ± standard error of the mean at each direction. See text for details.

At the pitch of maximum radiance, neither the mean brightness (Fig. 8A, Table 1, Test A2, Swoop dir.), hue angle (Fig. 8B, Table 1, Test A3, Swoop dir.), nor saturation (Fig. 8C, Table 1, Test A4, Swoop dir.) of the radiance varied significantly with the orientation of the swoop direction relative to the solar azimuth. Also, for brightness and saturation there was no difference between the male groups (Figs. 8A and B) how it changed with swoop bearing (Table 1, Tests A2, A3, and A4, Swoop dir.*Group) in the value of the color parameter (Table 1, Tests A2, A3, and A4, Group). The male groups did not differ in how hue angle changed with swoop bearing (Table 1, Test A4, Swoop dir.*Group), but mated males had a consistently and significantly higher hue angle (Table 1, Test A4, Group), that is, they were bluer, as was expected from the difference between the group in the hue of their reflectance.

**Fig. 8.**
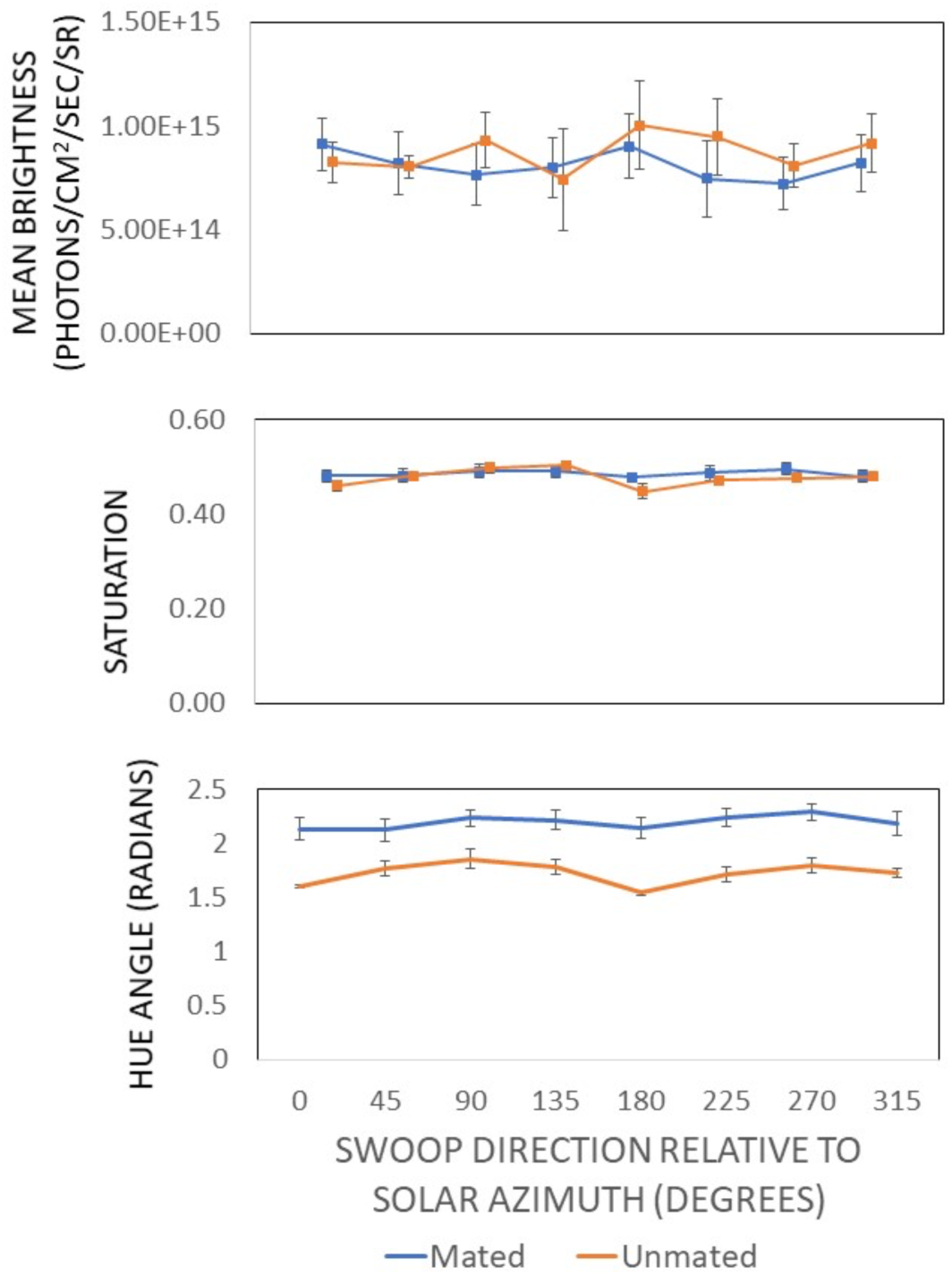
Color parameters of the male’s radiance at the pitch of maximum brightness as a function of the swoop bearing relative to the solar azimuth. Six male butterflies were sampled. The error bars indicate the mean ± standard error of the mean at each direction.

We then examined how the properties of the radiance from the dorsal hindwing surface visible to a female during a swoop changed as the male’s wings passed through the pitch that produced the maximum radiance visible to the female. Specifically, we looked at how relative brightness, hue angle and saturation changed every 10° starting 30° before the pitch of maximum radiance and ending at 30° after the pitch of maximum radiance (Fig. 9). The mean brightness increased greatly and significantly as the wing approached the pitch of maximum radiance and then declined quickly as the pitch went beyond the pitch of maximum brightness (Fig. 9A, Table 1, Test B1). The hue angle and saturation of the visible radiance also changed slightly but significantly with wing pitch (Fig. 9B and C, Tests B2 and B3). The change in saturation with wing pitch paralleled that of brightness with the brightest radiances being most saturated. Hue angle increased, that is, became more blue with wing pitch. Coupling these results with the observation that the male moves from below the female to directly in front of her in about 100 msec (Fig. 3) suggests that the female will see a brief, bright flash of saturated blue radiance from the male’s wings with a duration on the order of 10’s of msec.

**Fig. 9.**
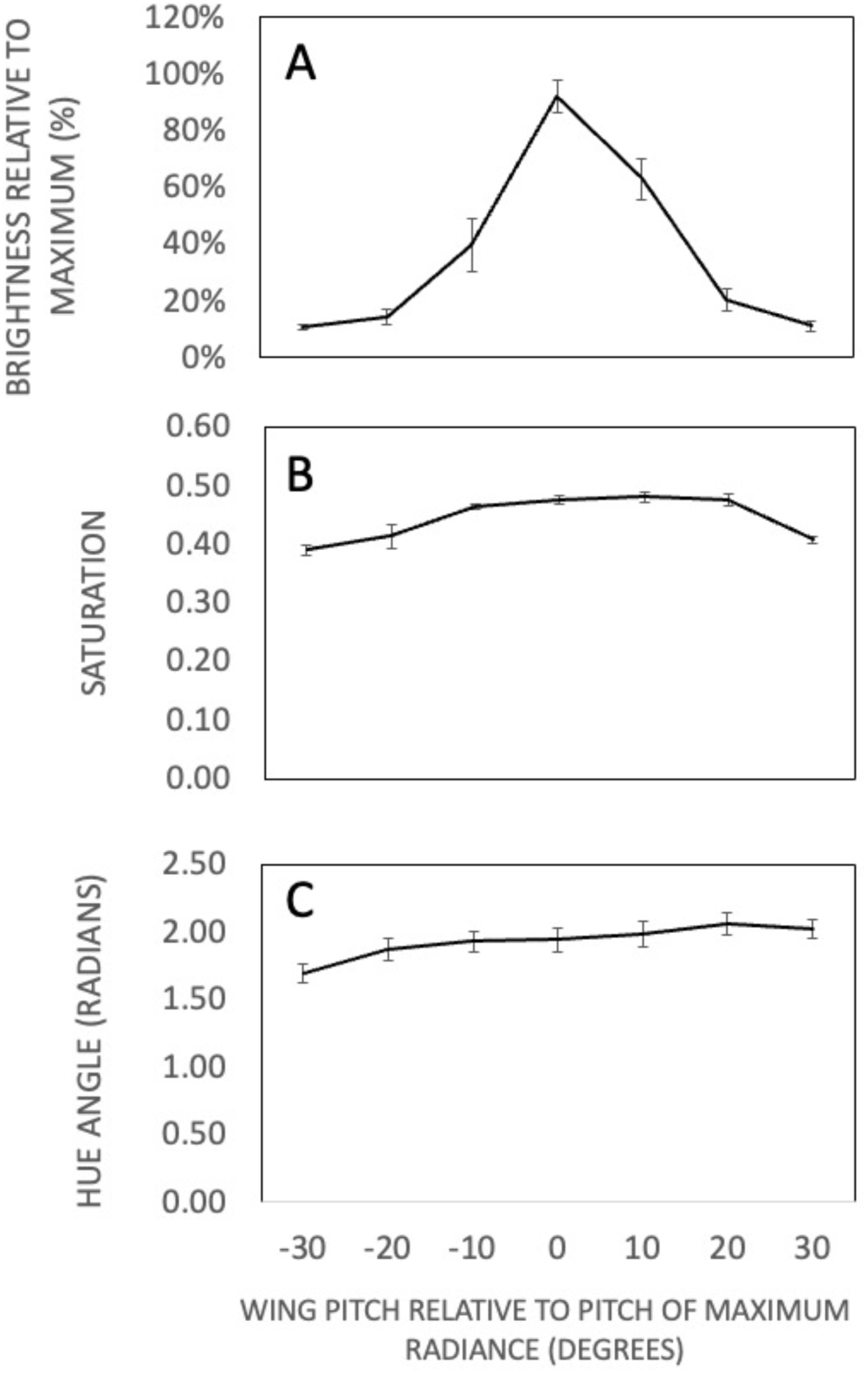
The brightness(A), saturation(B) and hue(C) of a *B. philenor* male’s coloration change during a swoop, as seen from the female’s position. Data points show the mean and standard deviation for four male butterflies sampled. See text for details.

## Discussion

Our analyses show that males of *B. philenor* perform a movement repeatedly during courtship that brings him from below the female to directly in front of her on an upward path. During this movement, called a swoop, the pitch of the male’s wings increases from horizontal to perpendicular relative to the ground. Several swoops are an obligate part of courtship leading to copulation in which the female is flying when the male approaches. That is, no flying female mated with a courting male unless he performed an average of 9-10 swoops. The male’s flight path during a swoop makes the male’s blue iridescent sexual signal appear very bright and saturated to the female at some point during a swoop regardless of the bearing of his flight path relative to the sun. Our broad conclusion is that the swoop is a display that adaptively enhances the visibility of the male’s iridescent color signal to the female during courtship and thereby enhancing his mating success.

### The swoop and signal delivery

Several features of the swoop appear to help overcome potential problems of signal transmission arising from the directionality of the male’s iridescent color signal. First, the swoop presents the dorsal hindwing surface at a variety of angles from near horizontal to near vertical in the female’s visual field. The visual field of *B. philenor* is expected to be large as it is in other butterflies (Rutowski et al. 2008, Bergman and Rutowski 2016) which means that the male’s dorsal hindwing radiance should be visible to the female whether he is directly below or directly in front of her. It is notable that these are the regions in the space around the female toward which the eye regions of smallest interommatidial angles (i.e. highest spatial resolution) in the eye of *B. philenor* are directed (Bergman and Rutowski 2016). The form of the swoop may have been driven by the position of this region of high acuity in the female’s visual field.

Second, regardless of the male’s orientation relative to the sun the swoop will include an angle that presents the male’s blue iridescence at maximum brightness and saturation to the female. The hue of the radiance visible to the female changes little as the male passes in front of her during a swoop. This is counter-intuitive given that iridescence is defined as a change in hue with angle of view (Simon 1971). The reason is that, in iridescence, hue changes with angle of illumination when coupled with a reciprocal change in the position of the viewer. The female’s angle of view of the male’s hindwing surface does not change substantially from 90° during the swoop. However, the source of the relevant incident light will change from blue sky to direct solar radiation.

Prior studies of *B. philenor* suggest that females attend to male blue coloration in their selection of mates. Rutowski and Rajyaguru (2013) provide experimental evidence that male saturation and, perhaps brightness, were positively correlated with male mating success. Sasaki et al. (2015) in a field study of the correlation between male coloration and recent mating success found that males with the bluest hues were most likely to have recently mated as indicated by the state of their reproductive tract. In our results, the female’s perception of brightness change dramatically as the male’s position changes during a swoop. The relative constancy of the hue and saturation of the male’s coloration with changes in wing pitch and orientation to the sun may make them better features to use in male assessment, however, evaluating this argument will require much more detailed information on exactly what features are most stimulating to females.

Males of *B. philenor* possess other sex-specific and potential signal-producing structures on their wings. The inner margin of each hindwing is folded dorsally over onto itself, forming an anal fold, in which scales with unique features are found (Miller 1987, Racheli and Oliverio 1993). These folds have been interpreted as scent-producing organs. At this point, no studies have examined whether special scents are being produced by these structures or that there are chemical signals involved in mate choice by females in this species. However, the experimental color modifications done by Rutowski and Rajyaguru (2013) did not alter the hindwing anal folds but still influenced male mating success. This suggests that visual signals predominate in female mate selection. Regardless, we cannot at this time completely discount the hypothesis that the swoop performed by males is also involved in transmitting a chemical signal to the female during courtship.

### Male courtship displays in butterflies

The swoop of *B. philenor* is a new addition to the diversity of male courtship behavior that has been reported in butterflies. Table S1 summarizes published reports that describe male behavior during courtship leading to copulation in 37 species of butterflies. In some species, the males do little other than approach a female and, once she alights, perch alongside her and attempt to couple. In other species, males have complex and stereotyped displays that they perform before attempting to couple. The diversity of form in these displays is high and includes a variety of species-specific body, wing and antennal movements and postures. Moreover, the diversity is high within, as well as, among clades. If taxonomy is taken as a surrogate for phylogeny, this suggests that these displays have evolved relatively quickly in the butterfly lineage. This is especially apparent within both the pierids and nymphalids. In both groups there is diversity in whether males perform distinctive displays and, among those that do perform displays, high diversity in the form of the displays. There is no clear phylogenetic signal among the displays although the sampling among taxonomic groups is uneven with few studies from the hesperids. Nonetheless, this speedy diversification is indicative of strong selection and is also evident in the evolution of coloration in the pierids (Kemp et al. 2005).

The swoops performed by male *B. philenor* are perhaps most like those described for *Argynnis paphia*. However, the shared features of this behavior seem most likely a result of convergence rather than homology. This is indicated especially by the high diversity of male courtship behavior in the nymphalids. However, the paucity of courtship studies in the swallowtails suggests more studies are needed of this group before we can be certain of the evolutionary history of swooping evolved within this group.

There has been a long history of interest in butterfly courtship dating back to Darwin (1871) who suggested that courtship in butterflies is a “prolonged affair.” The data reported in Table S1 suggest that, in fact, courtship leading to copulation in butterflies is stunningly brief, in some cases only a few seconds and generally less than a minute. We suspect that Darwin was basing his statement on observations of interactions between males and unreceptive females that did not end in copulation. Reports of unsuccessful courtships in the published reports often include cases in which courtship lasted for 10 min or more. Of special interest is that in species in which females are known to be selecting among conspecific males (*Eurema hecabe* (Kemp 2008), *Colias eurytheme* (Papke et al. 2007), *B. philenor* (Rutowski and Rajaguru 2013), successful courtship durations are on the order of seconds, not minutes. Apparently, females assess males with surprising speed.

Female preference is a strong candidate for the relevant selection pressure producing this diversity. In fact, several of the displays like the swoop have been interpreted as behavioral adaptations to enhance the delivery of signals that are important in female choice, e.g. the delivery of chemical signals via hairpencilling by males in the Queen (*D. gilippus*, Brower et al. 1965) and via wing snaps in the Gulf Fritillary (*Agraulis vanillae*, Rauser and Rutowski 2003; Rutowski and Schaefer 1984), and visual signals in the Common Eggfly (*H. bolina*, White et al. 2015).

Table S1 includes species, in addition to *B philenor*, in which females use iridescent male coloration in mate selection. Females in *H. bolina* choose males based on the dorsal spots of iridescent color on their wings (Kemp 2008). Interestingly, males in this species also perform a display that enhances the delivery of their visual signal (White et al. 2015), but the display is very different in form from that of *B. philenor*. Female sulphur butterflies also prefer to mate with males with a colorful iridescence on their dorsal wing surfaces (Papke et al., 2007, Kemp 2008), but in these species the male’s courtship behavior does not suggest any special features that enhance the visibility of the iridescent signal to females. Further studies of the pierid courtship will hopefully reveal just how and when during courtship females assess male iridescent coloration.

Among the species for which distinctive displays have been described, there are three listed in Table S1, each in a different family (*Nathalis iole, Agraulis vanilla, B. philenor*), in which females in some circumstances have mated with males who have not performed the described display. At least in some circumstances the display is not a requisite feature of courtship leading to copulation. If these displays are a product of sexual selection, it is not clear why, in some contexts, females accept males who do not perform them. One possibility is that females vary in their receptivity such that some have very low thresholds for accepting a mating.

### Lessons for the study of iridescent signals

Our experiences with the design and implementation of the measurements made during this study suggest several strategies that may prove valuable in other studies of the delivery of iridescence color signals. First, we affirm Fleishman et al.’s caveat (2006) that measuring radiances directly avoids several potential errors and complexities of inferring radiance from the product of reflectance and irradiance. An alternative is to measure perceived radiances using photographic techniques as Simpson and McGraw (2019b) have done for hummingbirds. However, this requires knowing the spectral sensitivities of the photoreceptors in the eye, information we do not have for *B. philenor*, and is not available for many species of interest. Second, it is important to describe carefully the spatial arrangement of sender, receiver, and light source and use that information to design measurements of radiance that accurately represent what is going on in nature and, therefore, how iridescent signals will be perceived by intended receivers. Third, color signal directionality does not necessarily mean there will be special displays (e.g., sulphurs and UV, no courtship displays), although even without obvious displays the question remains as to how the signal is detected by the female. We encourage students of iridescent color signals to keep these thoughts in mind.

## Supporting information

Information for supplemental video

Courtship video clips

## Acknowledgements

Kaci Fankhauser, Brittany Williams, and Ashley Lewis assisted with the analysis of courtship video and Michael Shillingburg helped with determination of swoop bearings. Dr. Thomas Cronin provided helpful guidance in the design and calculation of the radiance measurements. The Desert Botanical Garden graciously allowed us to use the Maxine and Jonathan Marshall Butterfly Pavilion for observations and video recording of courtship. This work was funded by an NSF grant (IOS 1145054) to RLR. NL was supported while in the lab of RLR at Arizona State University by a National Science Foundation Graduate Research Fellowship (DGE 0802261). For all this help we are very grateful.

