## Supplemental Table 1 for "Male behavior in a swallowtail butterfly (*Battus philenor*) ensures directional iridescent sexual signal is visible to females during courtship"

### Male Courtship Behavior in butterflies

Table S1 summarizes the courtship descriptions found in the literature as of 2022 using Google Scholar and other resources for 37 species of butterflies. For a published description to be included in this table, it had to include observations of multiple courtships leading to copulation. In most instances, these observations were made on male interactions with virgin females either in flight cages or in the field. Most reports also included information on the duration of these interactions, although in some cases, only rough estimates of the duration of successful courtship were offered. Selected summary information and statements on courtship durations from the published accounts are offered in the table.

Table S1 also notes for each species whether males perform a distinctive display as a prelude to copulation. We listed a display as any behavior reported by the authors that is beyond the male approaching, chasing, and hovering over the female before landing and attempting to mate with the female.

Table S1. Species for which courtship behavior has been described from observations of male-female interactions leading to copulation. In some species, distinctive displays were observed in some successful courtship but not all. Those species are indicated by "±" in the "Male display" column. .

| Family | | Species | Source | Male display | Courtship Duration |
|---|---|---|---|---|---|
| Pieridae | | | | | |
| | Pierinae | | | | |
| | | *Pieris protodice* | Rutowski 1979 | none | Median, female initially flying: 7 sec |
| | | *Pieris rapae* | Imafuku et al. 2021 | none | Mean: 9.2 sec |
| | | *Anthocaris cardamines* | Wiklund and Forsberg 1985 | none | Mean: 20.9 sec |
| | Coliadinae | | | | |
| | | *Eurema lisa* | Rutowski 1978 | none | Mean, with female flutter response: 4 sec |
| | | *Eurema daira* | Rutowski 1983 | wing waving | Mean, one display bout: 4 sec |
| | | *Eurema mandarina* | Imafuku et al. 2021 | none | Mean: 6.2 sec |
| | | *Nathalis iole* | Rutowski 1981 | wing spread (±) | Mean, with display: 14 sec |
| | | *Phoebis sennae* | Rutowski 1983 | none | Mean: 7 sec |
| | | *Colias eurytheme* | Silberglied and Taylor 1978 | none | "generally less than 5 sec" (pg. 218) |
| | | | Rutowski 1985 | none | Median, female initially flying: 9 sec |
| | Dismorphinae | | | | |

| | | | |
|---|---|---|---|
| *Leptidea synapsis* | Friberg et al. 2008; Wiklund 1977 | proboscis waving | Mean: 41 sec; range: 15-180 sec |
| *Leptidea reali* | Friberg et al. 2008 | proboscis waving | Mean: 127 sec |
| Nymphalidae | | | |
| Danainae | | | |
| *Danaus gilippus* | Brower et al. 1965 | hairpencilling | Mean: 55.2 sec |
| *Danaus plexippus* | Pliske 1975 | aerial takedown (±) | Mean: 70.9 sec |
| Apaturninae | | | |
| *Asterocampa leilia* | Rutowski and Gilchrist 1988 | none | Not reported |
| Limenitidinae | | | |
| *Limentis camilla* | Lederer 1960 | none | Not reported |
| Nymphalinae | | | |
| *Euphydryas chalcedona* | Rutowski and Gilchrist 1987 | none | Mean: 77.1 sec |
| | | wing clap on substrate | |
| *Hypolimnas bolina* | Rutowski 1992; White et al. 2014 | quivering flight | Not reported |
| Heliconiinae | | | |
| *Agraulis vanillae* | Rutowski and Schaefer 1984 | wing clap (±) | Mean, with display: 11 sec |
| *Heliconius erato* | Klein and de Araújo 2010 | androconia exposition | Range: 5-92 sec |
| *Argyreus hyperbius* | Imafuku et al. 2021 | none | Mean: 70.3 |
| *Argynnis paphia* | Magnus 1950 | swooping in flight, | Not reported |

| | | | |
|---|---|---|---|
| Satyrinae | | | |
| *Ypthima argus* | Imafuku et al. 2021 | bowing | Mean: 21.4 sec |
| *Hipparchia semele* | Tinbergen et al. 1942 | bowing | Not reported |
| *Bicyclus anynana* | Nieberding et al. 2008 | flickering, thrusting | “average courtship duration only a few sec” |
| Papilionidae | | | |
| Papilioninae | | | |
| *Battus philenor* | Rutowski et al., this study | swoop (±) | Mean with swoops: 17.2 sec |
| *Papilio glaucus* | Krebs 1988 | none | Mean: 58 sec |
| Lycaenidae | | | |
| Lycaeninae | | | |
| *Heodes virgaurae* | Douwes 1976 | none | Not reported |
| *Lycaena argyrognomon* | Lundgren and Bergström 1975 | none | Not reported |
| *Lycaena phlaeas* | Imafuku et al. 2021 | none | Mean: 203.8 sec |
| Theclinae | | | |
| *Artopoetes pryeri* | Imafuku et al. 2021 | none | Mean: 10.7 sec |
| *Japonica lutea* | Imafuku et al. 2021 | none | Mean: 25.6 sec |
| *Shirozua jomasi* | Imafuku et al. 2021 | none | Mean: 18.8 sec |
| *Neozephyrus japonicus* | Imafuku et al. 2021 | none | Mean: 90.8 sec |
| *Chrysozephyrus* | | | |

| | | | |
|---|---|---|---|
| *smaragdinus* | Imafuku et al. 2021 | none | Mean: 159.5 sec |
| Polyommatinae | | | |
| *Zizeeria maha* | Imafuku et al. 2021 | none | Mean: 42.6 sec |
| *Lampides boeticus* | Imafuku et al. 2021 | none | Mean: 167.6 sec |
| Hesperidae | | | |
| Hesperiinae | | | |
| *Hylephila phylaeus* | Shapiro 1975 | head-wing behavior | Median: <40 sec |
