## Supplementary material for "Male behavior in a swallowtail butterfly (*Battus philenor*) ensures directional iridescent sexual signal is visible to females during courtship": Information for supplemental video

Videos of swoops

The video contains three clips of *Battus philenor* courtship filmed in the Maxine and Jonathan Marshall Butterfly Pavilion at the Desert Botanical Gardens, Phoenix, Arizona. The clips were filmed at 240 frames per sec and so are displayed at one-eighth normal speed. Clip 1 shows five swoops directed by male at a female with marks on her ventral hindwing made with white correction fluid. Clip 2 shows two swoops. Clip 3 shows seven swoops performed by a male with white marks (correction fluid) on his right dorsal forewing and left ventral hindwing.
